# A protective and broadly binding antibody class engages the influenza virus hemagglutinin head at its stem interface

**DOI:** 10.1101/2023.12.13.571543

**Authors:** Holly C. Simmons, Joel Finney, Ryutaro Kotaki, Yu Adachi, Annie Park Moseman, Akiko Watanabe, Shengli Song, Lindsey R. Robinson-McCarthy, Valerie Le Sage, Masayuki Kuraoka, E. Ashley Moseman, Garnett Kelsoe, Yoshimasa Takahashi, Kevin R. McCarthy

**Affiliations:** Center for Vaccine Research, University of Pittsburgh School of Medicine, Pittsburgh, PA, USA.; Department of Microbiology and Molecular Genetics, University of Pittsburgh School of Medicine, Pittsburgh, PA, USA; Department of Integrative Immunobiology, Duke University, Durham, North Carolina, USA; Department of Immunology, National Institute of Infectious Diseases, Tokyo 162-8640, Japan; Department of Surgery, Duke University, Durham, North Carolina 27710, USA

## Abstract

Influenza infection and vaccination impart strain-specific immunity that protects against neither seasonal antigenic variants nor the next pandemic. However, antibodies directed to conserved sites can confer broad protection. Here we identify and characterize a class of human antibodies that engage a previously undescribed, conserved epitope on the influenza hemagglutinin (HA) protein. Prototype antibody S8V1-157 binds at the normally occluded interface between the HA head and stem. Antibodies to this HA head-stem interface epitope are non-neutralizing *in vitro* but protect against lethal influenza infection in mice. Antibody isotypes that direct clearance of infected cells enhance this protection. Head-stem interface antibodies bind to most influenza A serotypes and seasonal human variants, and are present at low frequencies in the memory B cell populations of multiple human donors. Vaccines designed to elicit these antibodies might contribute to “universal” influenza immunity.

## INTRODUCTION

Influenza pandemics arise from antigenically novel zoonotic influenza A viruses transmitted to humans from animals. Historically, pandemic viruses generally have become endemic and have continued to circulate as seasonal viruses. Sustained viral circulation is enabled by on-going antigenic evolution that leads to escape from population-level immunity elicited by previous exposures ^1^. Although antibodies provide the strongest protection against infection, they also drive the antigenic evolution of the viral surface proteins ^1^. Therefore, neither infection nor the seasonal flu vaccine confers enduring immunity against future seasonal variants or new pandemic viruses. Nevertheless, broadly protective monoclonal antibodies that engage conserved influenza sites have been isolated from human donors ^2^. Passive transfer of these antibodies to animal models imparts broadly protective immunity. A next generation influenza vaccine that elicits similar antibodies would likely confer more durable protection than that offered by current seasonal vaccines ^3–5^.

The influenza hemagglutinin (HA) protein is the major target of protective antibodies ^6^. HA facilitates cell entry by attaching to cells via an interaction with its receptor, sialic acid, and by acting as a virus-cell membrane fusogen. HA is synthesized as a polyprotein, HA0, which forms homotrimers that are incapable of undergoing the full series of conformational rearrangements required for membrane fusion. Cellular proteases (often resident on the target cell) cleave the HA0 into HA1 and HA2 domains, resulting in a fusion-competent trimer ^7–9^. HA1 includes the globular HA head that contains the receptor binding site (RBS), while HA2 contains the helical stem regions that rearrange during endosomal acidification to drive fusion of viral and cellular membranes. The requirement for HA to transit through multiple conformations during fusion likely accounts for the intrinsic propensity of HA0 or HA1-HA2 to transiently explore states that deviate from its prefusion conformation, which reproducibly forms ordered protein crystal lattices that yield high-resolution structures determined by X-ray crystallography^10–13^.

Large genetic and antigenic differences separate influenza A HA serotypes, which are classified into group 1 (H1, H2, H5, H6, H8, H9, H11, H12, H13, H16) and group 2 (H3, H4, H7, H10, H14, H15). Serotypes of a second glycoprotein, neuraminidase (NA), are denoted by a similar convention; consequently, influenza viruses are named by their HA and NA content (e.g., H2N2, H10N8). Currently, divergent H1N1 and H3N2 viruses circulate as seasonal human influenza viruses. Humans can produce antibodies that engage surfaces conserved between serotypes and thereby provide cross-serotype protection. These epitopic regions include the HA RBS, stem, anchor, head interface, the so-called “long alpha-helix”, and the HA2 β-hairpin ^2,14–19^.

Here, we report a class of human antibodies directed to a previously unreported, widely conserved site at the base, or neck, of the HA head. In the “resting state” prefusion structure of the HA trimer, the epitope for prototype antibody S8V1-157 is occluded at the interface of the head and stem. Biochemical, cellular, and *in vivo* passive transfer experiments indicate that this HA head-stem interface epitope is sufficiently exposed to allow antibodies to bind and confer strong protection against lethal influenza virus infection in murine challenge models. Many humans harbor antibodies recognizing this epitope; some of these antibodies bind HAs from divergent influenza serotypes, groups, and >50 years of human seasonal virus antigenic variation. Immunogens that elicit such antibodies might be included in influenza vaccines intended to confer broader protection.

## RESULTS

### A class of antibodies engages the interface of the HA head and stem regions

By culturing individual human memory B (Bmem) cells and then screening the culture supernatants containing secreted IgGs, we identified (as reported previously ^20^) 449 HA-reactive antibodies from four donors (S1, S5, S8 and S9). These antibodies represent the Bmem cells circulating in the blood at the time the donors were immunized with the TIV 2015-2016 seasonal influenza vaccine (visit 1; V1), or 7 days later (visit 2; V2). From donors S1 and S8, we found four IgGs (S1V2-17, S1V2-60, S1V2-65 and S8V1-157) that had a novel pattern of HA reactivity (Figure 1A). All bound seasonal H1s and H3s and an H5 HA; they also bound recombinant HA constructs comprising only the HA head, indicating the IgGs did not target stem epitopes (Figure 1A). These four antibodies competed with each other for HA-binding, implying overlapping epitopes; however, the four antibodies did not compete with other well-characterized antibodies that engage conserved, structure-verified, broadly protective epitopes spanning the HA molecule (Figure 1B).

**Figure 1:**
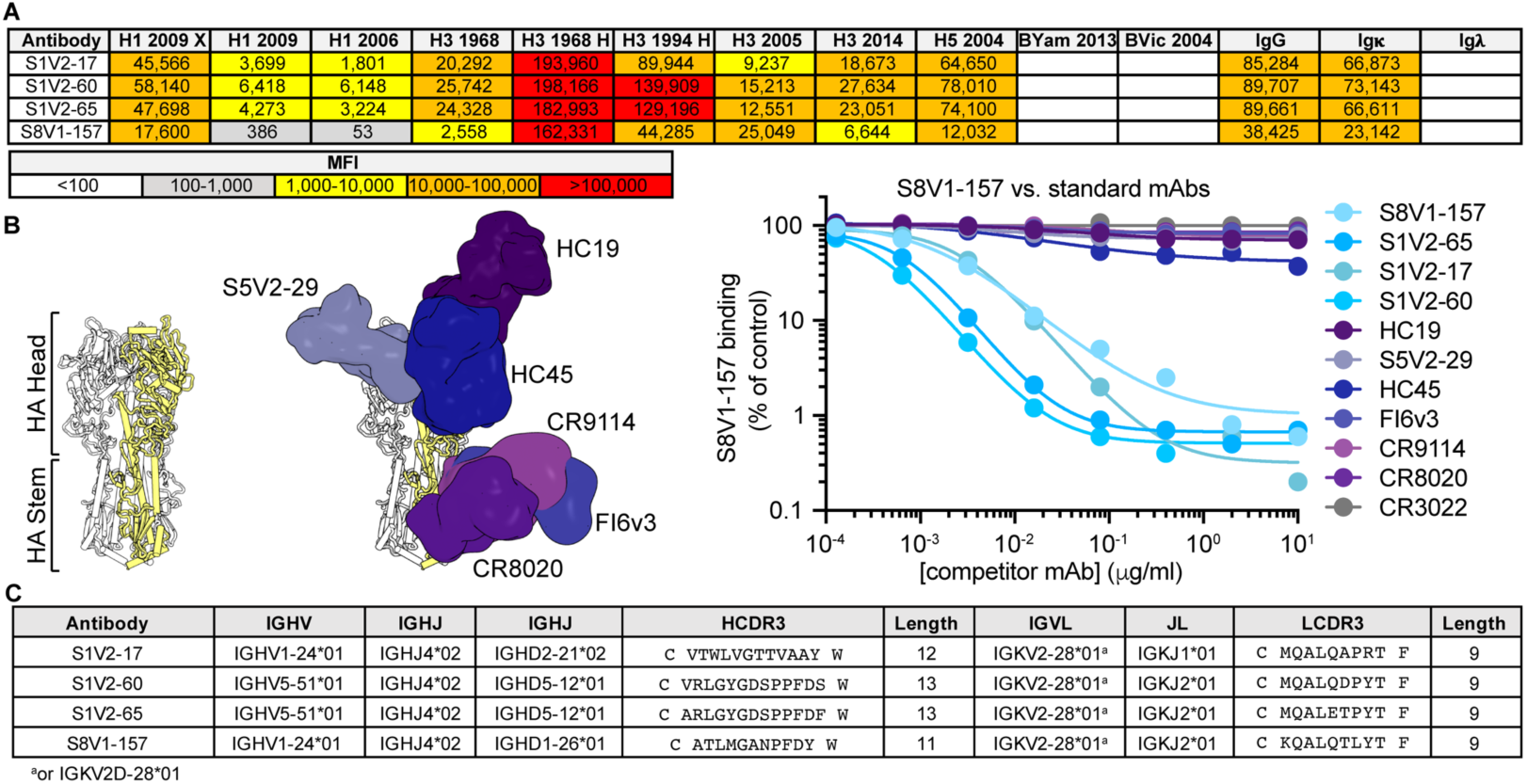
Identification of a novel, broadly binding, HA head-directed antibody class. A. Luminex screening of Bmem-cell Nojima culture supernatants identified four antibodies that broadly react with influenza A HA FLsEs and heads. The mean fluorescence intensity (MFI) values are colored according to the key. B. In a Luminex competitive binding assay, the four antibodies from (A) that share a pattern of reactivity did not compete with antibodies that engage known HA epitopes, but compete with each other for HA binding. Structures of Fab-HA complexes were aligned on an HA trimer from A/American black duck/New Brunswick/00464/2010(H4N6) (PDB: 5XL2)^21^. Fab structures include HC19^32^ (PDB 2VIR), S5V2-29^20^ (PDB 6E4X), HC45^57^ (PDB 1QFU), CR9114^58^ (PDB 4FQY), CR8020^59^ (PDB 3SDY) and FI6v3^47^ (PDB 3ZTJ). SARS-CoV antibody CR3022^33^ was used as an HA non-binding control. C. The cross-competing HA antibodies share genetic signatures.

All four new antibodies have light chains encoded by IGKV2-28*01 or IGKV2D-28*01 (Figure 1C), with identical germline coding sequences. The heavy chains are encoded by either IGHV1-24*01 or IGHV5-51*01 and have short (11-13 amino acid) third complementarity-determining regions (HCDR3s). S1V2-60 and S1V2-65 are clonally related, while S1V2-17 and S8V1-157 share a common IGHV1-24*01-IGKV2-28*01 pairing, despite their derivation from different donors.

We determined the structure of S8V1-157 complexed with a monomeric HA head construct. Crystals were only obtained using an A/American black duck/New Brunswick/00464/2010(H4N6) (H4-NB-2010) HA (Figure 2A and Table S1). S8V1-157 engages an epitope at the base of the head, just where it faces the stem ^21^. In the context of trimeric, full-length, prefusion HA, this epitope is occluded by the association of HA1 with HA2 (Figure 2A). Therefore, displacement of the HA1 head from the HA2 helical stem would be required for B cells or antibodies to recognize this site.

**Figure 2:**
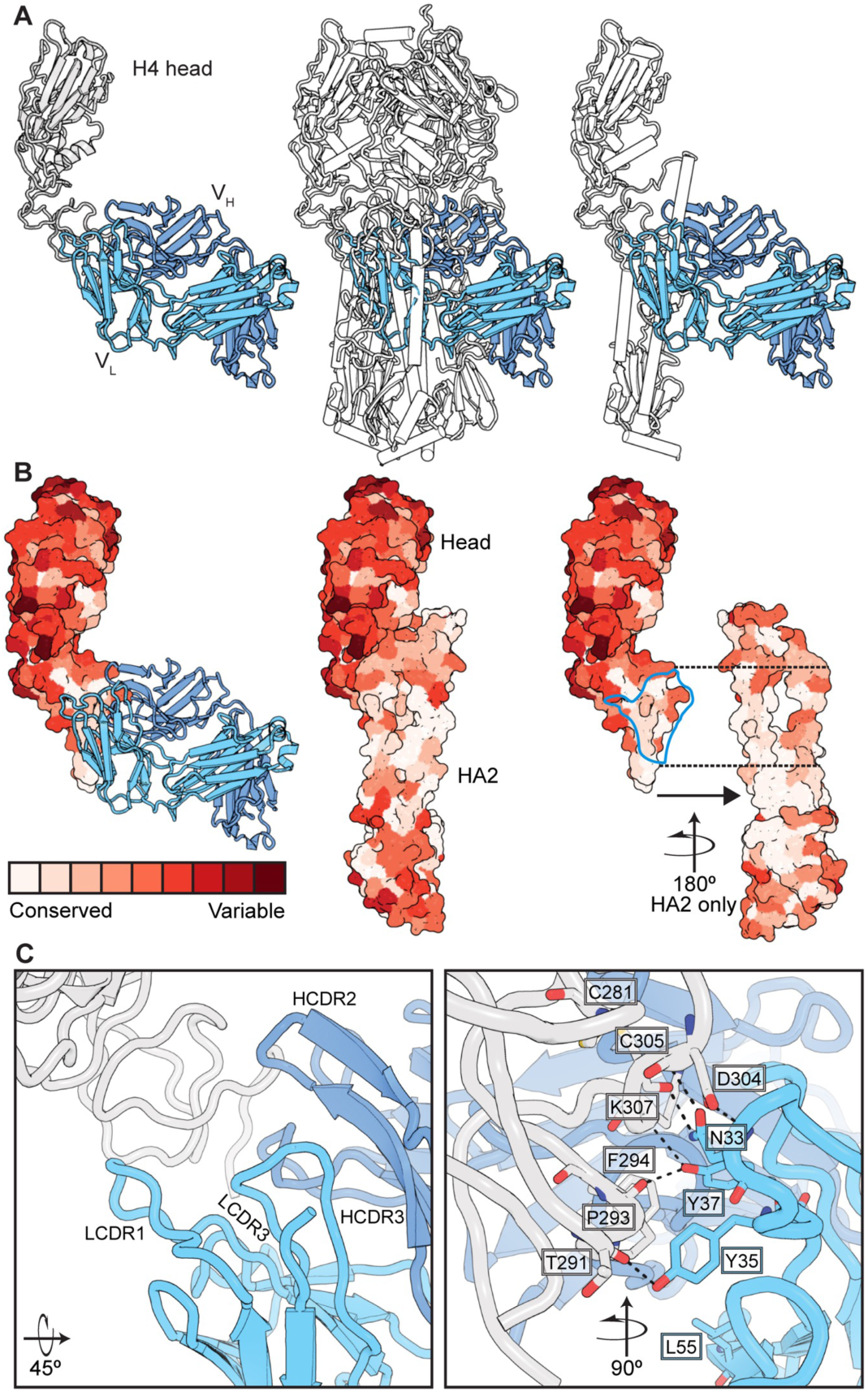
Human antibodies engage a recessed surface at the head-stem interface of the influenza HA molecule. A. Structure of antibody S8V1-157 complexed with the HA head of A/American black duck/New Brunswick/00464/2010(H4N6) colored in gray. The heavy chain is colored darker blue and the light chain is lighter blue. Engagement of this site is incompatible with the defined prefusion H4 HA trimer^21^, colored in white (PDB: 5XL2) or with individual HA monomers. B. A surface projection showing the degree of amino acid conservation among HAs engaged by this antibody class (see Figure 3). The head-stem epitope is circumscribed in blue in the rightmost panel. Conservation scores were produced using ConSurf ^60,61^. C. Key S8V1-157 contacts. The orientation relative to panel A is indicated.

S8V1-157 contacts residues mediating the intra-protomer interaction, which are conserved across divergent HAs (Figures 2B and S1). Evolutionary constraints imposed by the requirement that HA1 and HA2 stably associate at neutral pH likely account for the conservation of the head-stem interface epitope, and by extension, the binding breadth of the corresponding antibodies.

The compact HCDR3 of S8V1-157 packs against the heavy chain CDR and framework regions (FR), HCDR1-FR1 and FR2-HCDR2, to produce a cleft between the heavy and light chains that accommodates the head-stem interface epitope (Figure 2C). Light chain CDR1 residues N33, Y35 and Y37 make extensive contacts with a conserved HA surface. The δ-amide of N33 donates and receives hydrogen bonds from the main chain of HA residue HA-C305. Its main chain amide also donates a hydrogen bond to the carboxyl group of HA-D304. Y35 donates a hydrogen bond to the main chain carbonyl of HA-T291 and participates in van der Waals interactions by stacking upon HA-P293. Y37 receives a hydrogen bond from the main chain residue HA-K307, donates a hydrogen bond to HA-P293, and participates in van der Waals contacts with the aryl group of HA-F294 and the aliphatic side chain of HA-K307.

The structure of the S8V1-157-HA complex explains the genetic features common to this class of head-stem interface antibodies. The LCDR1 NXYXY motif is germline-encoded by a small subset of IGKV-genes. Of these, only IGKV2-28/IGKV2D-28a also encode a hydrophobic residue at position 55. L55 participates in van der Waals contacts with HA-P293 (Figure 2C) and contributes to a hydrophobic surface on S8V1-157 that complements a hydrophobic patch within its epitope on the HA head (Figure S2). Polar residues at position 55 in otherwise similar light chains are predicted to be less favorable in this local environment. Similar sequence features are not present in the IGLV locus. Additionally, the short HCDR3 (common among these antibodies) creates shallow cavity that accommodates the hydrophobic surfaces that would normally pack against HA2 in the prefusion HA trimer.

### Antibodies to the HA head-stem interface epitope are broadly binding

We determined the binding of head-stem interface antibodies to divergent HA serotypes using recombinantly expressed soluble HA ectodomains and recombinantly expressed antibodies in enzyme-linked immunosorbent assays (ELISA). All HAs except H5 and H7 were expressed as HA0, which is incapable of transitioning to the postfusion conformation without first being processed by a trypsin-like protease^8,22^. H5 and H7 HAs contain polybasic cleavage sites that are processed by endogenous furin-like proteases in the expressing cell line, resulting in HA1-HA2^23–28^.

The HA head-stem interface antibodies had very similar breadth: each bound all seasonal H1s (1977-2019) and H3s (1968-2020) assayed, and also bound several other HA subtypes within groups 1 and 2 (Figures 3A and S3). Typically, if an HA was bound by one head-stem epitope antibody, it was also bound by the other three (Figure 3). Affinities and breadth of binding were generally higher for group 2 HAs but extended to group 1 HAs, including seasonal H1, pandemic H2, and pre-pandemic H5 HAs. Overall, these antibodies engaged HAs from 11 of the 16 non-bat influenza A HA serotypes. Failure to engage specific HAs was not due to epitope inaccessibility, since the antibodies also did not bind matched, soluble, HA head that present the HA head-stem epitope without steric hindrance (Figures S4 and S5). None of the head-stem interface antibodies bound a truncated HA head lacking the S8V1-157 epitope. All four antibodies likely engage a common, discrete epitope comprising the terminus of the HA head.

**Figure 3:**
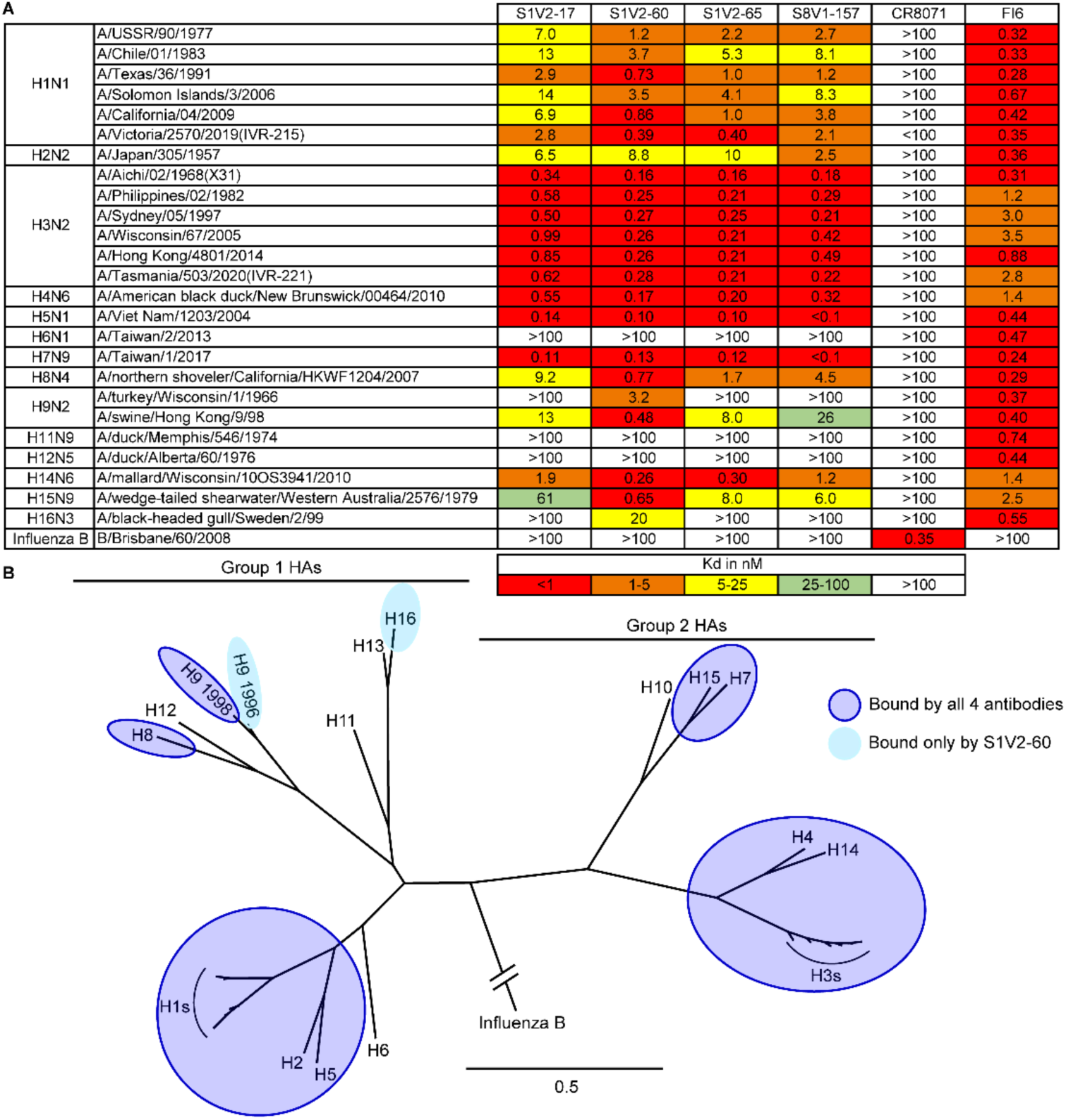
Breadth of HA binding by HA head-stem epitope antibodies. A. Equilibrium dissociation constants (K_d_), determined by ELISA. Broadly binding influenza A HA antibody FI6v3 ^47^ and influenza B HA antibody CR8071 ^58^ served as binding controls. B. Phylogenetic relationships of HAs used in our panel. HAs bound by HA head-stem epitope antibodies are indicated. Binding data from Figure S4 are incorporated into panel B.

In a complementary flow cytometry assay (in the absence of exogenous trypsin^19,29^), HA head-stem epitope antibodies bound divergent HAs stably expressed in their native, transmembrane form on the cell surface (Figure 4). The binding pattern was consistent with our ELISA data. We used a curated panel of antibodies (largely overlapping with those in Figure 1), including K03.12^30^ and H5.3^31^ which recognize the fully exposed receptor binding site. HA head-stem interface antibodies generally labeled cells less brightly than did control antibodies directed to solvent-exposed epitopes on the HA head (e.g., K03.12, H5.3, HC19); however, head-stem interface antibodies labeled cells comparably to the HA head-interface antibody S5V2-29 ^20^. Together, these observations suggest that HA interface epitopes are not occupied as fully by antibody as are fully exposed epitopes. Epitope accessibility and/or differences in the number of IgG molecules bound per HA trimer may account for these differences.

**Figure 4:**
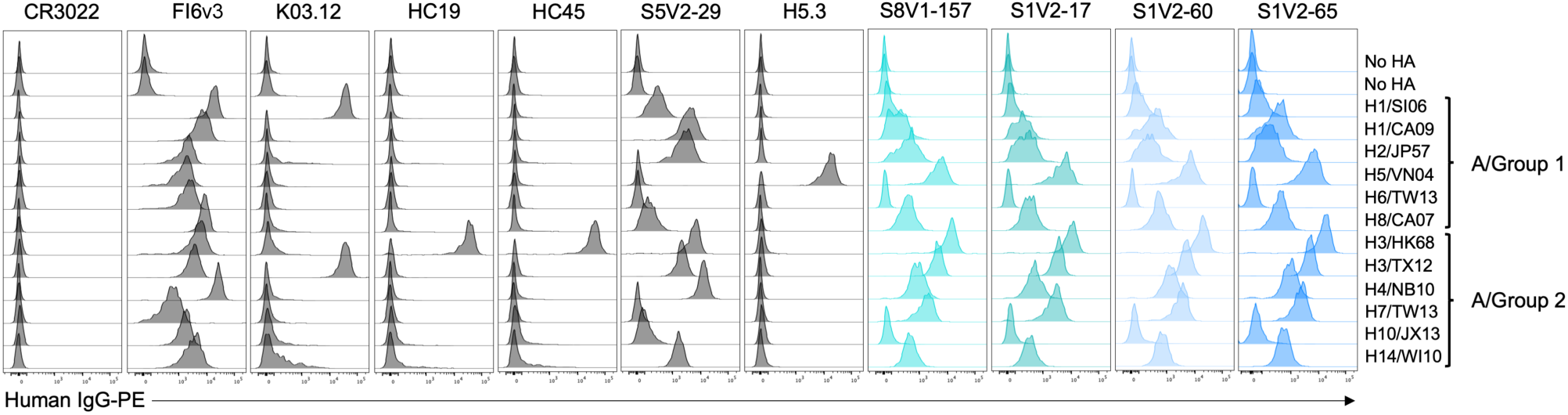
HA head-stem epitope antibodies bind cell surface-anchored HA. Flow cytometry histograms depict the fluorescence intensities of recombinant IgG binding to K530 cell lines expressing recombinant, native HA on the cell surface. K530 cells were labeled with 400 ng/ml of the four head-stem epitope antibodies or control antibodies targeting the HA receptor binding site (HC19^32^, K03.12^30^ and H5.3^31^), the head interface (S5V2-29^20^), a lateral head epitope (HC45)^57^, stem (FI6v3^47^), or SARS-CoV spike protein (CR3022^33^). HA abbreviations correspond to: H1/SI06: A/Solomon Islands/3/2006(H1N1), H1/CA09: A/California/04/2009(H1N1), H2/JP57: A/Japan/305/1957(H2N2), H5/VN04: A/Viet Nam/1203/2004(H5N1), H6/TW13: A/Taiwan/2/2013(H6N1), H8/CA07: A/northern shoveler/California/HKWF1204/2007(H8N4),H3/HK68: A/Aichi/02/1968(H3N2)(X31), H3/TX12: A/Texas/50/2012(H3N2), H4/NB10:A/American black duck/New Brunswick/00464/2010, H7/TW13: A/Taiwan/1/2017(H7N9), H10/JX13: A/Jiangxi/IPB13/2013(H10N8),H14/WI10: A/mallard/Wisconsin/10OS3941/2010(H14N6).

To determine whether head-stem interface antibodies can engage native HA0 on the cell surface, 293F cells were transiently transfected with an expression vector encoding full-length, membrane-anchored A/Aichi/02/1968(H3N2)(X-31) HA. Western blotting analysis using an antibody against a linear epitope contained within HA1 confirmed that the transfected cells expressed only unprocessed, 83 kD HA0 (Fig. S5B). Processed HA1, which under reducing and denaturing conditions would be observed as a 55 kD band, was not detected. Flow cytometry analysis showed that S8V1-157 labeled the transfected cells (Fig. S5C), as did antibodies recognizing the RBS (HC19) and head interface epitope (S5V2-29). The head-stem interface epitope is therefore exposed, if only transiently, on native HA0 expressed on the cell surface.

### Antibodies to the HA head-stem epitope protect against lethal influenza virus infection

S8V1-157 failed to neutralize influenza virus *in vitro*, in an assay in which virus and antibody were co-incubated, added to cells, and neutralization scored four days post-infection (Figure S6). To determine if head-stem interface antibodies confer protection *in vivo*, we produced the antibodies as recombinant mouse IgG1 or IgG2c, then passively transferred the antibodies to mice and challenged them with a lethal dose of influenza virus. In mice, the IgG2c isotype potently directs protective, Fc-dependent, effector functions, including antibody-dependent cellular cytotoxicity (ADCC) and complement deposition (ADCD). In contrast, the IgG1 isotype does not appreciably stimulate these effector functions. Each mouse received 150 μg (∼7.5 mg/kg) of the head-stem interface antibodies S8V1-157 or S5V2-65, HC19^32^ (which potently neutralizes A/Aichi/02/1968(H3N2)(X-31)), S5V2-29^20^ (an HA-head interface antibody that protects against lethal infection by Fc-dependent mechanisms), or CR3022^33^ (SARS-CoV antibody). The ∼7.5 mg/kg antibody dose is comparable or below commonly administered doses of antiviral monoclonal antibody therapeutics^33–38^.

HC19 protected mice from infection-induced weight loss, while the irrelevant antibody CR3022 offered no protection, requiring recipient mice to be ethically euthanized by day 10 post-infection (Figure 5A). All animals administered HA head-stem interface antibodies S8V1-157 or S1V2-65 recovered after experiencing mild to moderate weight loss; by day 10 post-infection, their body weights were similar to the HC19-treated group. IgG2c versions of S8V1-157 and S1V2-65 conferred slightly better protection than the corresponding IgG1s, indicating that Fc effector functions had a role in controlling the infection (Figure 5B-D). Similarly, head interface antibody S5V2-29 protected better against infection-induced weight loss when infused as an IgG2c. Thus, despite failing to neutralize the initial infection, HA head-stem interface antibodies protected influenza-infected mice against severe disease and mortality.

**Figure 5:**
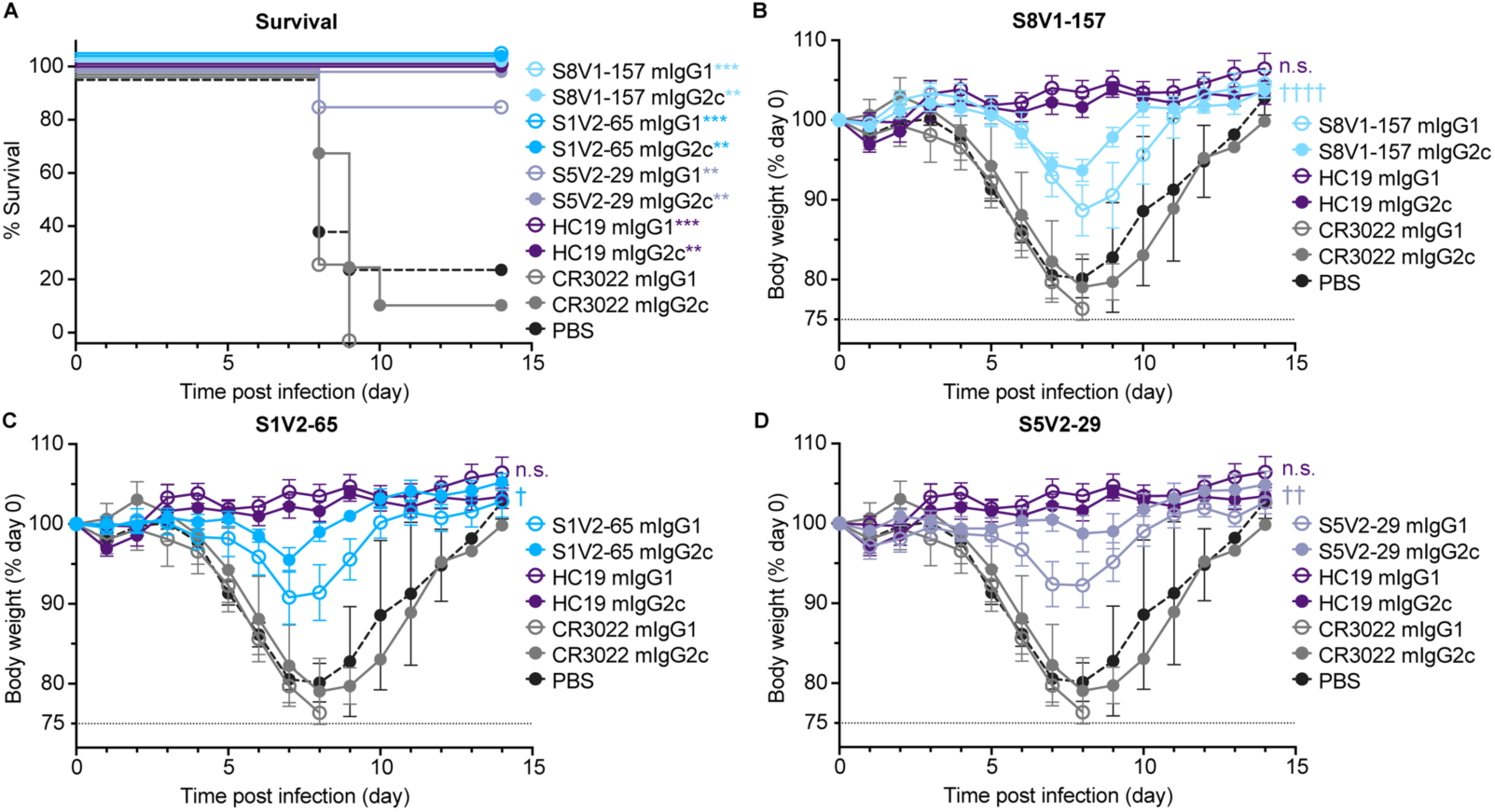
HA head-stem epitope antibodies protect against lethal influenza virus infection and severe disease. C57BL/6 mice (n = 7 per group) were intraperitoneally injected with 150 μg of recombinant antibody via intraperitoneal injection three hours prior to intranasal challenge with 5xLD50 of A/Aichi/02/1968(H3N2)(X31). Mice were weighed daily and euthanized at a humane endpoint of 25% loss of body weight. Antibodies passively transferred included musinized IgG1 and IgG2c versions of HA head-stem epitope antibodies S8V1-157 and S1V2-65, neutralizing antibody HC19^32^, head interface antibody S5V2-29^20^ and SARS-CoV antibody CR3022^33^. Mice injected with PBS were included as an additional control. A. Post-infection survival rate. B-D. Body weight curves for infected mice administered S8V1-157 (B), S1V2-65 (C), or S5V2-29 (D) antibodies, compared with controls. *p < 0.05 and ***p < 0.001 compared with isotype control CR3022. Not significant (n.s.), p 0.05; † p < 0.05, †† p < 0.01, and †††† p < 0.0001 IgG2c compared with IgG1.

### The HA head-stem epitope is immunogenic in humans

We screened additional human Bmem cell cultures for S8V1-157-competing antibodies (Figure 6). The samples were taken from seven donors, including the same subjects as before (S1, S5, S8, S9), but vaccinated and sampled during subsequent flu seasons (see Materials and Methods for details); subject S12, who was immunized and sampled at the same times; and subjects KEL01 and KEL03, who were sampled after receiving TIV in 2014-2015. From 528 clonal cultures that produced HA-binding IgG, we identified eight additional supernatants that inhibited S8V1-157 binding to HA by >90%. (Figure 6A). In a screen of antibody reactivity, five of the eight competing antibodies reacted with both group 1 and group 2 HAs.

**Figure 6:**
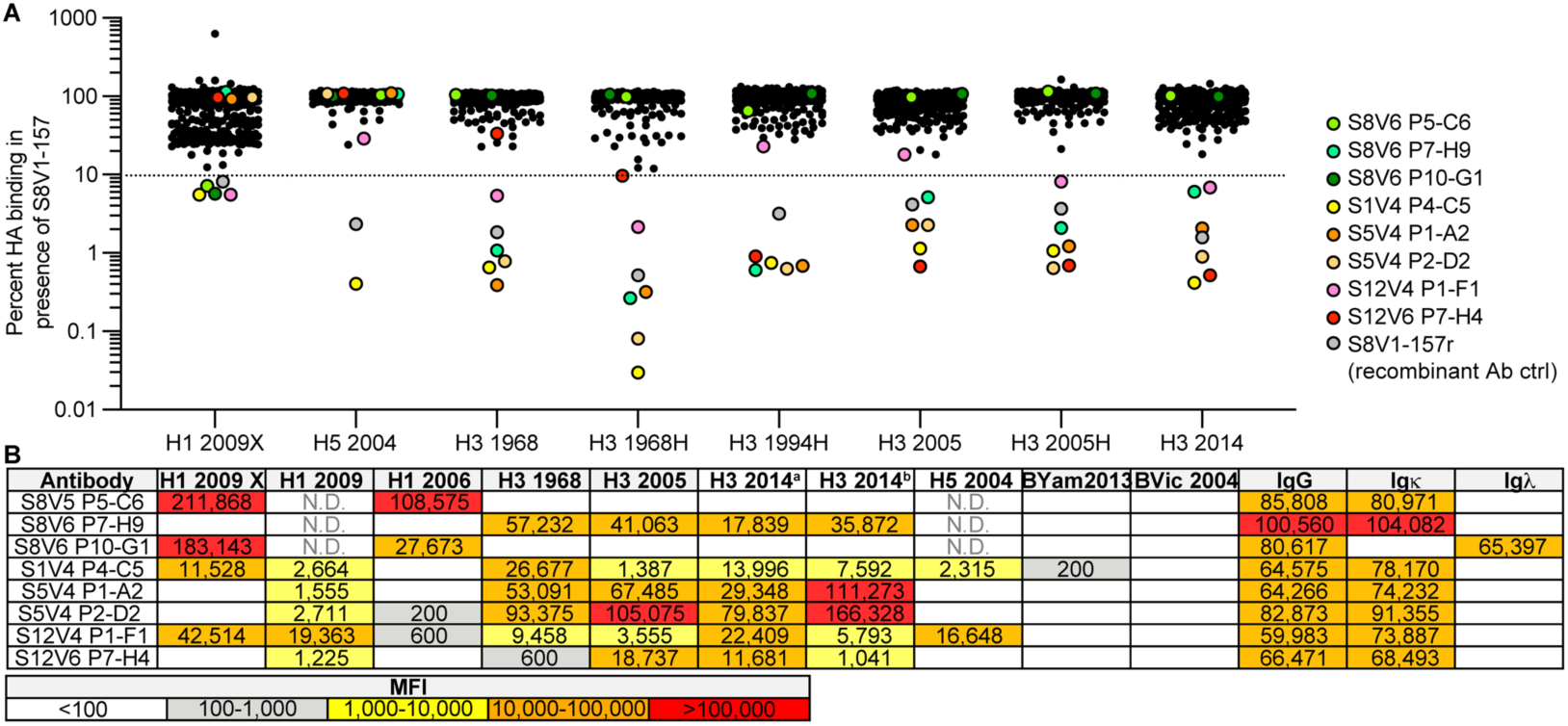
The HA head-stem epitope epitope is immunogenic in humans. A. An additional 528 Nojima culture supernatants from donors K01, K03, S1, S5, S8, S9 and S12 were screened for competition with a recombinant musinized S8V1-157 IgG1 for HA binding. Culture supernatants that inhibited S8V1-157 binding by >90% are colored and specified. B. HA reactivity of S8V1-157-competing Nojima culture supernatants, as determined by multiplex Luminex assay. N.D.: not determined.

Paired heavy and light chain sequences were recovered from seven of the eight S8V1-157-competing antibodies (Figure S7). Two antibodies, S1V4-P4-C5 and S12V6-P7-H4, use an IGKV2-28 light chain paired with IGHV1-24*01, like S8V1-157 and S1V2-17. All four antibodies have short, 11-12 amino acid HCDR3s. These features, shared with S8V1-157, likely define a public antibody class present in at least three human donors. The other newly identified S8V1-157 competitors use other *IGHV* and *IGKV* genes and have varying HCDR3 lengths (10-19 amino acids). These antibodies could either have footprints that overlap the head-stem interface epitope or that prevent its exposure.

## DISCUSSION

We identified and characterized a human antibody response directed to a previously unrecognized, widely conserved, cryptic epitope on the influenza HA head. These antibodies bind broadly to divergent HA serotypes found in animals and seasonal antigenic variants of human viruses. Prophylactic passive transfer of these antibodies to mice protects against lethal influenza virus disease. Approximately 1% of circulating, HA-reactive Bmem recognize the head-stem interface and are present in multiple donors both before and after seasonal influenza vaccination, though most such Bmem were isolated following vaccination. Head-stem interface antibodies have a bias, but not a restriction, for IGKV2-28 usage. Across the 12 known examples we find no additional constraints. Other humans are therefore likely to produce similar antibodies and to have the capacity to mount polyclonal, broadly protective antibody responses.

That HA head-stem interface antibodies confer robust protection against lethal influenza challenge demonstrates that the epitope is exposed on infected cells and/or virions, if only transiently. Exposure of the HA head-stem interface would be predicted to require a large-scale displacement of the HA head from HA2, and additional conformational rearrangements within the HA trimer might accompany such events. Nevertheless, our *in vitro* experiments, which used recombinant HA0 (either as soluble ectodomain or native, membrane-anchored HA on the surface of transiently-transfected cells), confirmed that head-stem interface antibodies must be able to recognize their epitope on some conformation of prefusion HA, because HA0 cannot transition to a stable postfusion structure^8^. Structural, biophysical, and computational approaches indicate that HA trimers transiently adopt conformations that expose epitopes normally occluded in the defined prefusion state^10–13^. Such transient fluctuations might explain how B cells and antibodies recognize ordinarily occluded epitopes^16,17,20,39–41^. In addition to binding HA0, head-stem interface antibodies might also directly recognize postfusion HA on the surface of infected cells, as has been proposed for some HA antibodies, including LAH31^17^, S1V2-72^19^ and m826^41^. It is plausible that, *in vivo*, HA0 is cleaved into HA1/HA2 by proteases and that some fraction of postfusion HA containing the exposed head-stem interface epitope accumulates on infected cell-surfaces and/or among the HAs presented to B cells in germinal centers.

Humans and mice mount antibody responses to several apparently occluded HA epitopes ^14–17,19,20,39–43^. Prevalence of such antibodies in immune repertoires indicates that these epitopes are immunogenic. That several of the human head-stem interface antibodies reported here had 5-11% IGHV gene mutation frequencies implies that the epitope was encountered repeatedly over recurrent influenza exposures (whether by infection or vaccination). The immunogenicity of the HA head-stem epitope poses a critical question for vaccinology: how does the antigen presented to a B cell in a germinal center reaction relate to its form on infectious virions or to what has been defined in the laboratory in considerable biochemical and structural detail? The existence of antibodies to interface/buried epitopes on other, unrelated viral glycoproteins demonstrates that our lack of understanding extends beyond HA^44^.

Several of the most conserved epitopes on HA are occluded. Adjacent protomers and/or membranes hinder access to some stem/anchor epitopes, and interface epitopes are normally hidden ^2,14,15,19,45^. Nevertheless, antibodies recognizing these epitopes protect against lethal influenza virus disease in small animal models. These antibodies are typically less potently neutralizing than antibodies targeting fully exposed sites, and their protection is often enhanced by or fully dependent on Fc-mediated effector functions ^16,17,19,20,40,43,46,47^. Their resistance to human influenza antigenic evolution and ability to engage emerging, pre-pandemic, and other animal viruses make their selective elicitation a potential strategy to improve current influenza vaccines. Given the characteristics of head-stem interface antibodies, a rigorous understanding of their potential to enhance human protective immunity and/or ameliorate disease is needed.

## METHODS

### Human subjects

Peripheral blood mononuclear cells (PBMCs) were obtained from human donors KEL01 (male, age 39) and KEL03 (female, age 39), under Duke Institutional Review Board committee guidelines. Written informed consent was obtained from both subjects. KEL01 and KEL03 received the trivalent inactivated seasonal influenza vaccine (TIV) 2014-2015 Fluvirin, which contained A/Christchurch/16/2010, NIB-74 (H1N1), A/Texas/50/2012, NYMC X-223 (H3N2), and B/Massachusetts/2/2012, NYMC BX-51B. Blood was drawn on day 14 post-vaccination, and PBMCs isolated by centrifugation over Ficoll density gradients (SepMate-50 tubes, StemCell Tech) were frozen and kept in liquid nitrogen until use.

PBMCs were also obtained from human donors S1 (female, age 51-55), S5 (male, age 21-25), S8 (female, age 26-30), S9 (female, age 51-55), and S12 (male, age 35-40), under Boston University Institutional Review Board committee guidelines. Written informed consent was obtained from all five subjects. Donors met all of the following inclusion criteria: between 18 and 65 years of age; in good health, as determined by vital signs [heart rate (<100 bpm), blood pressure (systolic ≤ 140 mm Hg and ≥ 90 mm Hg, diastolic ≤ 90 mm Hg), oral temperature (<100.0 °F)] and medical history to ensure existing medical diagnoses/conditions are not clinically significant; can understand and comply with study procedures, and; provided written informed consent prior to initiation of the study. Exclusion criteria included: 1) life-threating allergies, including an allergy to eggs; 2) have ever had a severe reaction after influenza vaccination; 3) a history of Guillain-Barre Syndrome; 4) a history of receiving immunoglobulin or other blood product within the 3 months prior to vaccination in this study; 5) received an experimental agent (vaccine, drug, biologic, device, blood product, or medication) within 1 month prior to vaccination in this study or expect to receive an experimental agent during this study; 6) have received any live licensed vaccines within 4 weeks or inactivated licensed vaccines within 2 weeks prior to the vaccination in this study or plan receipt of such vaccines within 2 weeks following the vaccination; 7) have an acute or chronic medical condition that might render vaccination unsafe, or interfere with the evaluation of humoral responses (includes, but is not limited to, known cardiac disease, chronic liver disease, significant renal disease, unstable or progressive neurological disorders, diabetes mellitus, autoimmune disorders and transplant recipients); 8) have an acute illness, including an oral temperature greater than 99.9°F, within 1 week of vaccination; 9) active HIV, hepatitis B, or hepatitis C infection; 10) a history of alcohol or drug abuse in the last 5 years; 11) a history of a coagulation disorder or receiving medications that affect coagulation. Subjects S1, S5, S8, S9, and S12 received seasonal influenza vaccination during three consecutive North American flu seasons (2015-2016, 2016-2017, 2017-2018), and had blood drawn on day 0 (pre-vaccination; visits 1, 3, and 5) and day 7 (post-vaccination, visits 2, 4, and 6) each year. During the 2015-2016 season (visits 1 and 2), the subjects received the TIV Fluvirin, which contained A/reassortant/NYMC X-181 (California/07/2009 x NYMC X-157) (H1N1), A/South Australia/55/2014 IVR-175 (H3N2), and B/Phuket/3073/2013. During the 2016-2017 season (visits 3 and 4), the subjects received the quadrivalent inactivated vaccine Flucelvax, containing A/Brisbane/10/2010 (H1N1), A/Hong Kong /4801/2014 (H3N2), B/Utah/9/2014, and B/Hong Kong/259/2010. During the 2017-2018 season (visits 5 and 6), the subjects received the quadrivalent inactivated vaccine Flucelvax, containing A/Singapore/GP1908/2015 IVR-180 (H1N1), A/Singapore/GP2050/2015 (H3N2), B/Utah/9/2014, and B/Hong Kong/259/2010.

### Cell lines

Human 293F cells were maintained at 37°C with 5-8% CO_2_ in FreeStyle 293 Expression Medium (ThermoFisher) supplemented with penicillin and streptomycin. HA-expressing K530 cell lines ^29^ (*Homo sapiens*) were cultured at 37°C with 5% CO_2_ in RPMI-1640 medium plus 10% FBS (Cytiva), 2-mercaptoethanol (55 μM; Gibco), penicillin, streptomycin, HEPES (10 mM; Gibco), sodium pyruvate (1 mM; Gibco), and MEM nonessential amino acids (Gibco). Madin-Darby canine kidney (MDCK) were maintained in Minimum Essential medium supplemented with 10% fetal bovine serum, 5 mM L-glutamine and 5 mM penicillin/streptomycin.

### Recombinant Fab expression and purification

The heavy and light chain variable domain genes for Fabs were cloned into a modified pVRC8400 expression vector, as previously described^48–50^. Fab fragments used in crystallization were produced with a noncleavable 6xhistidine (6xHis) tag on the heavy chain C-terminus. Fab fragments were produced by polyethylenimine (PEI) facilitated, transient transfection of 293F cells. Transfection complexes were prepared in Opti-MEM (Gibco) and added to cells. 5 days post transfection, cell supernatants were harvested and clarified by low-speed centrifugation. Fabs were purified by passage over TALON Metal Affinity Resin (Takara) followed by gel filtration chromatography on Superdex 200 (GE Healthcare) in 10 mM tris(hydroxymethyl)aminomethane (tris), 150 mM NaCl at pH 7.5 (buffer A).

### Single B cell Nojima cultures

Nojima cultures were previously performed^18–20^. Briefly, peripheral blood mononuclear cells (PBMCs) were obtained from four human subjects S1 (female, age range 51-55), S5 (male, age 21-25), S8 (female, age 26-30), and S9 (female, age 51-55). Single human Bmem cells were directly sorted into each well of 96-well plates and cultured with MS40L-low feeder cells in RPMI1640 (Invitrogen) containing 10% HyClone FBS (Thermo scientific), 2-mercaptoethanol (55 μM), penicillin (100 units/ml), streptomycin (100 μg/ml), HEPES (10 mM), sodium pyruvate (1 mM), and MEM nonessential amino acid (1X; all Invitrogen). Exogenous recombinant human IL-2 (50 ng/ml), IL-4 (10 ng/ml), IL-21 (10 ng/ml) and BAFF (10 ng/ml; all Peprotech) were added to cultures. Cultures were maintained at 37° degrees Celsius with 5% CO_2_. Half of the culture medium was replaced twice weekly with fresh medium (with fresh cytokines). Rearranged *V(D)J* gene sequences for human Bmem cells from single-cell cultures were obtained as described^20,48,51^. Specificity of clonal IgG antibodies in culture supernatants and of rIgG antibodies were determined in a multiplex bead Luminex assay (Luminex Corp.). Culture supernatants and rIgGs were serially diluted in 1 × PBS containing 1% BSA, 0.05% NaN_3_ and 0.05% Tween20 (assay buffer) with 1% milk and incubated for 2 hours at room temperature with the mixture of antigen-coupled microsphere beads in 96-well filter bottom plates (Millipore). After washing three times with assay buffer, beads were incubated for 1 hour at room temperature with Phycoerythrin-conjugated goat anti-human IgG antibodies (Southern Biotech). After three washes, the beads were re-suspended in assay buffer and the plates read on a Bio-Plex 3D Suspension Array System (Bio-Rad).

### Recombinant HA expression and purification

Recombinant HA head constructs and Full-length HA ectodomains (FLsE) were expressed by polyethylenimine (PEI) facilitated, transient transfection of 293F cells. To clone HA heads, synthetic DNA for the region was subcloned into a pVRC8400 vector encoding a C-terminal rhinovirus 3C protease site and a 6xHis tag. To produce FLsE constructs, synthetic DNA was subcloned into a pVRC8400 vector encoding a T4 fibritin (foldon) trimerization tag and a 6xHis tag. Transfection complexes were prepared in Opti-MEM (Gibco) and added to cells. 5 days post transfection, cell supernatants were harvested and clarified by low-speed centrifugation. HA was purified by passage over TALON Metal Affinity Resin (Takara) followed by gel filtration chromatography on Superdex 200 (GE Healthcare) in buffer A. HA heads used for crystallography underwent the following additional purification steps. HA heads were cleaved using the Pierce 3C HRV Protease Solution Kit (Ref 88947) and passed over TALON Metal Affinity Resin to capture cleaved tags. Cleaved HA heads were then further purified by gel filtration chromatography on Superdex 200 (GE Healthcare) in buffer A.

### ELISA

Five hundred nanograms of rHA FLsE or HA head were adhered to high-capacity binding, 96 well-plates (Corning 9018) overnight in PBS pH 7.4 at 4°C. HA coated plates were washed with a PBS-Tween-20 (0.05%v/v) buffer (PBS-T) and then blocked with PBS-T containing 2% bovine serum albumin (BSA) for 1 hour at room temperature. Blocking solution was then removed, and 5-fold dilutions of IgGs (in blocking solution) were added to wells. Plates were then incubated for 1 hour at room temperature. Primary IgG solution was removed and plates were washed three times with PBS-T. Secondary antibody, anti-human IgG-HRP (Abcam ab97225) diluted 1:10,000 in blocking solution, was added to wells and incubated for 30 minutes at room temperature. Plates were then washed three times with PBS-T. Plates were developed using 150μl 1-Step ABTS substrate (ThermoFisher, Prod#37615). Following a brief incubation at room temperature, HRP reactions were stopped by the addition of 100μl of 1% sodium dodecyl sulfate (SDS) solution. Plates were read on a Molecular Devices SpectraMax 340PC384 Microplate Reader at 405 nm. KD values for ELISA were obtained as follows. All measurements were performed in technical triplicate. The average background signal (no primary antibody) was subtracted from all absorbance values. Values from multiple plates were normalized to the FI6v3^47^ standard (FluA20 was used for the HA head assays) that was present on each ELISA plate. The average of the three measurements were then graphed using GraphPad Prism (v9.0). KD values were determined by applying a nonlinear fit (One site binding, hyperbola) to these data points. The constraint that Bmax must be greater than 0.1 absorbance units was applied to all KD analysis parameters.

### Recombinant IgG expression and purification

The heavy and light chain variable domains of selected antibodies were cloned into modified pVRC8400 expression vectors to produce full length human IgG1 heavy chains and human lambda or kappa light chains. IgGs were produced by transient transfection of 293F cells as specified above. Five days post-transfection supernatants were harvested, clarified by low-speed centrifugation, and incubated overnight with Protein A Agarose Resin (GoldBio) at 4°C. The resin was collected in a chromatography column and washed with one column volume of buffer A. IgGs were eluted in 0.1M Glycine (pH 2.5) which was immediately neutralized by 1M tris (pH 8.5). Antibodies were then dialyzed against phosphate buffered saline (PBS) pH 7.4.

### Recombinant mIgG expression and endotoxin free purification

The heavy and light chain variable domains of selected antibodies were cloned into respective modified pVRC8400 expression vector to produce full length murine IgG1 and IgG2C heavy chains and murine lambda or kappa light chains. IgGs were produced by transient transfection of 293F cells as specified above. Five days post-transfection supernatants were harvested, clarified by low-speed centrifugation. 20% volume of 1M 2-morpholin-4-ylethanesulfonic acid (MES) pH 5 and Immobilized Protein G resin (Thermo Scientific Prod#20397) were added to supernatant and sample was incubated overnight at 4°C. The resin was collected in a chromatography column and washed with one column volume of 10mM MES, 150 mM NaCl at pH 5. mIgGs were eluted in 0.1M Glycine (pH 2.5), which was immediately neutralized by 1M tris (pH 8.5). Antibodies were then dialyzed against phosphate buffered saline (PBS) pH 7.4. All mIgGs administered to mice were purified using buffers made with endotoxin-free water (HyPure Cell Culture Grade Water, Cytiva SH30529.03) and were dialyzed into endotoxin-free Dulbecco’s PBS (EMD Millipore, TMS-012-A). Following dialysis, endotoxins were removed using Pierce High Capacity Endotoxin Removal Spin Columns (Ref 88274).

### Virus microneutralization assays

Two-fold serial dilutions of 50 ug/mL of HC19, S8V1-157 or CR3022 were incubated with 10^3.3^ TCID_50_ of A/Aichi/02/1968 H3N2 (X-31) influenza virus for 1 hour at room temperature with continuous rocking. Media with TPCK was added to 96-well plates with confluent MDCK cells before the virus:antibody mixture was added and left on the cells for the duration of the experiment. After 4 days, cytopathic effect was determined and the neutralizing antibody titer was expressed as the reciprocal of the highest dilution of serum required to completely neutralize the infectivity each virus on MDCK cells. The concentration of antibody required to neutralize 100 TCID_50_ of virus was calculated based on the neutralizing titer dilution divided by the initial dilution factor, multiplied by the antibody concentration.

### Protection

C57BL/6 female and male mice were obtained from the Jackson Laboratory. All mice were housed under pathogen-free conditions at Duke University Animal Care Facility. Eight to 10-week old mice were injected *i.p.* with 150 μg of recombinant antibody diluted to 200 μl in PBS. Three hours later, mice were anesthetized by *i.p.* injection of ketamine (85 mg/kg) and xylazine (10 mg/kg) and infected intranasally with 3×LD_50_ (1.5×10^4^ PFU) of A/Aichi/2/1968 X-31 (H3N2) in 40 μL total volume (20 μL/nostril). Mice were monitored daily for survival and body weight loss until 14 days post-challenge. The humane endpoint was set at 20% body weight loss relative to the initial body weight at the time of infection. All animal experiments were approved by the standards and guidance set forth by Duke University IACUC.

Female C57BL/6J mice at 8 weeks old, purchased from Japan SLC, were i.p. injected with 150 μg of the antibodies in 150 μL PBS. Three hours later, the mice were intranasally infected with 5LD_50_ of A/Aichi/02/1968(H3N2)(X31), kindly gifted from Dr. Takeshi Tsubata (Tokyo Medical and Dental University), under anesthesia with medetomidine-midazolam-butorphanol. Survival and body weight were daily assessed for 14 days with a humane endpoint set as 25% weight loss from the initial body weight. The experimental procedures were approved by the Animal Ethics Committee of the National Institute of Infectious Diseases, Japan, and performed in accordance with the guidelines of the Institutional Animal Care and Use Committee. Statistical significance of body weight change and survival of mice after lethal influenza challenge was calculated by Two-way ANOVA test and Mantel-Cox test, respectively, using GraphPad Prism (v10.0) software

### Competitive inhibition assay

Competitive binding inhibition was determined by a Luminex assay, essentially as described^20,30^. Briefly, serially diluted human rIgGs or diluted (1:10) Bmem cell culture supernatants were incubated with HA-conjugated Luminex microspheres for 2 h at room temperature or overnight at 4°C. S8V1-157 mouse IgG1 was then added at a fixed concentration (100 ng/ml final for competition with rIgGs, or 10 ng/ml final for competition with culture supernatants) to each well, and incubated with the competitor Abs and HA-microspheres for 2 h at room temperature. After washing, bound S8V1-157 was detected by incubating the microspheres with 2 μg/ml PE-conjugated rat anti-mouse IgG1 (SB77e, SouthernBiotech) for 1 hr at room temperature, followed by washing and data collection. Irrelevant rIgG or culture supernatants containing HA-nonbinding IgG were used as non-inhibiting controls.

### Flow cytometry

Flow cytometry analysis of rIgG binding to HA-expressing K530 cell lines was performed essentially as described ^29^. Briefly, K530 cell lines were thawed from cryopreserved aliquots and expanded in culture for μ3 days. Pooled K530 cells were incubated at 4°C for 30 min with 0.4 μg/ml recombinant human IgGs diluted in PBS plus 2% fetal bovine serum. After washing, cells were labeled with 2 μg/ml PE-conjugated goat anti-human IgG (Southern Biotech) for 30 min at 4°C. Cells were then washed and analyzed with a BD FACSymphony A5 flow cytometer. Flow cytometry data were analyzed with FlowJo software (BD).

Flow cytometry analysis of rIgG binding to HA-expressing 293F cells was performed as follows. 293F cells were transfected using PEI with either plasmid encoding full-length HA from A/Aichi/02/1968(H3N2)(X31) or with empty vector. 36 hours post-transfection, cells were incubated at 4°C for one hour with 0.4 μg/ml recombinant human IgGs diluted in PBS plus 2% fetal bovine serum. After washing, cells were labeled with BB515 mouse anti-human IgG (BD Biosciences) for 30 min at 4°C at the manufacturer’s recommended concentration, followed by washing and fixation in 2% paraformaldehyde. Cells were analyzed with a BD LSRFortessa flow cytometer. Flow cytometry data were analyzed with FlowJo software (BD).

### Western blots

293F cells were transfected using PEI with either plasmid encoding full-length HA from A/Aichi/02/1968(H3N2)(X31) or with empty vector. 24 hours post-transfection, cells were lysed in RIPA buffer (25mM Tris pH7.6, 150mM NaCl, 1% NP40 alternative, 1% sodium deoxycholate, 0.1% SDS). Cell lysates were clarified of debris and boiled with Laemmli buffer with 2-mercaptoethanol. Samples were run on 4-20% acrylamide gels and transferred to nitrocellulose membranes. Membranes were blocked in 5% milk in PBS with 0.05% Tween 20 and probed with anti-HA tag antibody (Thermo-Fisher A01244-100; 0.3 μg/ml), followed by IR800-conjugated goat anti-mouse immunoglobulin (LiCor 926-32210; 0.1 μg/ml) and DyLight680-conjugated mouse anti-rabbit GAPDH antibody (BioRad MCA4739D680; 0.3 μg/ml). Membranes were imaged using a LiCor Odyssey CLx imager.

### Crystallization

S1V2-157 Fab fragments were co-concentrated with the HA-head of A/American black duck/New Brunswick/00464/2010(H4N6) at a molar ratio of ∼1:1.3 (Fab to HA-head) to a final concentration of ∼20 mg/ml. Crystals of Fab-head complexes were grown at 18°C in hanging drops over a reservoir solutions containing 0.1 M Lithium sulfate, 0.1 M Sodium chloride, 0.1 M 2-(N-morpholino)ethanesulfonic acid (MES) pH 6.5 and 30% (v/v) poly(ethylene glycol) (PEG) 400. Crystals were cryprotected with 30% (v/v) PEG 400, 0.12 M Lithium sulfate, 0.3 M Sodium chloride, and 0.06 M MES pH 6.5. Cryprotectant was added directly to the drop, crystals were harvested, and flash cooled in liquid nitrogen.

### Structure determination and refinement

We recorded diffraction data from a single moderate to poorly diffracting crystal at the Advanced Photon Source on beamline 24-ID-C. The dataset was strongly anisotropic, variable mosaicity (average 1.4°) and comprised contributions from multiple lattices. Efforts to improve these crystals were unsuccessful. Data were processed and scaled (XSCALE) with XDS^52^. Molecular replacement was carried out with PHASER^53^, dividing each complex into four search models (HA-head, Vh, Vl and constant domain). Search models were 5XL2, 6EIK, 5BK5 and 6E56.We carried out refinement calculations with PHENIX^54^ and model modifications, with COOT^55^. Refinement of atomic positions and B factors was followed by translation-liberation-screw (TLS) parameterization. All placed residues were supported by electron density maps and subsequent rounds of refinement. Final coordinates were validated with the MolProbity server^56^. Data collection and refinement statistics are in Table S1. Figures were made with PyMOL (Schrödinger, New York, NY.

## DATA AND SOFTWARE AVAILABILITY

Coordinates and diffraction data have been deposited at the PDB, accession number 8US0. Antibody sequences have been deposited in NCBI GenBank, accession numbers OR825693-OR825700.

## ACKNOWLEDGMENTS

We thank the many members of our Program Project Consortium for advice and discussion. X-ray diffraction data were recorded at beamline ID-24-C (operated by the Northeast Collaborative Access team: NE-CAT) at the Advanced Photon Source (APS, Argonne National Laboratory). We thank NE-CAT staff members for advice and assistance in data collection. NE-CAT is funded by NIH grant P30 GM124165. APS is operated for the DOE Office of Science by Argonne National Laboratory under contract DE-AC02-06CH11357. The research was supported by NIAID Program Project Grant P01 AI089618 (to G.H.K.), funds from the University of Pittsburgh Center for Vaccine Research (to K.R.M), and Japan Agency for Medical Research and Development grant JP22fk0108141 (to Y.A. and Y.T.)

## Supporting Figures

**Supporting Table 1:**
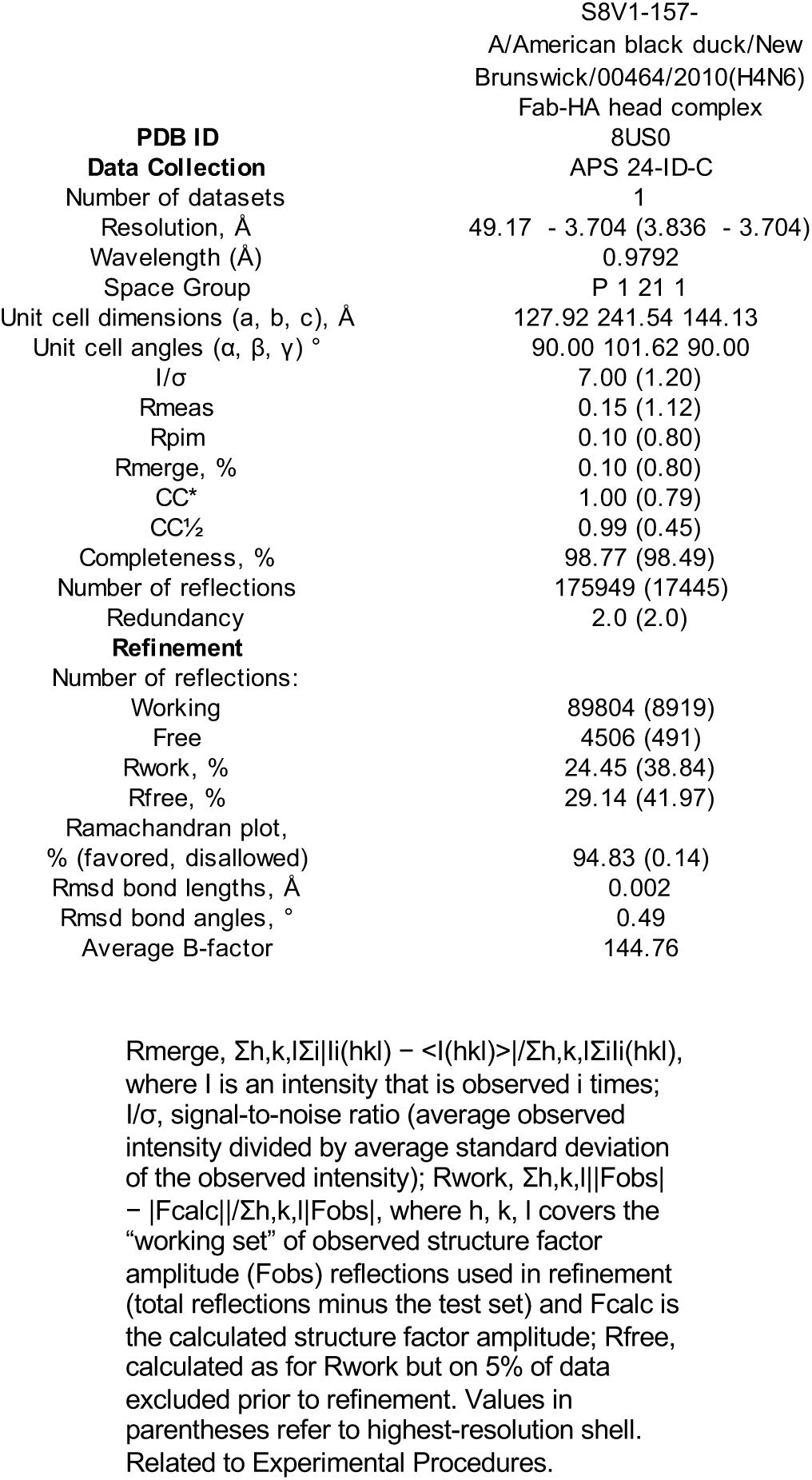
Data collection and refinement statistics.

|  |  |
| --- | --- |
|  | S8V1-157- |
|  | A/American black duck/New |
|  | Brunswick/00464/2010(H4N6) |
|  | Fab-HA head complex |
| **PDB ID** | 8US0 |
| **Data Collection** | APS 24-ID-C |
| Number of datasets | 1 |
| Resolution, Å | 49.17 - 3.704 (3.836 - 3.704) |
| Wavelength (Å) | 0.9792 |
| Space Group | P 1 21 1 |
| Unit cell dimensions (a, b, c), Å | 127.92 241.54 144.13 |
| Unit cell angles (α, β, γ) ° | 90.00 101.62 90.00 |
| I/σ | 7.00 (1.20) |
| Rmeas | 0.15 (1.12) |
| Rpim | 0.10 (0.80) |
| Rmerge, % | 0.10 (0.80) |
| CC* | 1.00 (0.79) |
| CC½ | 0.99 (0.45) |
| Completeness, % | 98.77 (98.49) |
| Number of reflections | 175949 (17445) |
| Redundancy | 2.0 (2.0) |
| **Refinement** |  |
| Number of reflections: |  |
| Working | 89804 (8919) |
| Free | 4506 (491) |
| Rwork, % | 24.45 (38.84) |
| Rfree, % | 29.14 (41.97) |
| Ramachandran plot,<br>% (favored, disallowed) | 94.83 (0.14) |
| Rmsd bond lengths, Å | 0.002 |
| Rmsd bond angles, ° | 0.49 |
| Average B-factor | 144.76 |

$R_{merge} = \frac{\sum_{h,k,l} \sum_i |I_i(hkl) - \langle I(hkl) \rangle|}{\sum_{h,k,l} \sum_i I_i(hkl)}$ , where I is an intensity that is observed i times; I/σ, signal-to-noise ratio (average observed intensity divided by average standard deviation of the observed intensity); Rwork, $\frac{\sum_{h,k,l} ||F_{obs}| - |F_{calc}||}{\sum_{h,k,l} |F_{obs}|}$ , where h, k, l covers the “working set” of observed structure factor amplitude (Fobs) reflections used in refinement (total reflections minus the test set) and Fcalc is the calculated structure factor amplitude; Rfree, calculated as for Rwork but on 5% of data excluded prior to refinement. Values in parentheses refer to highest-resolution shell. Related to Experimental Procedures.

**Figure S1:**
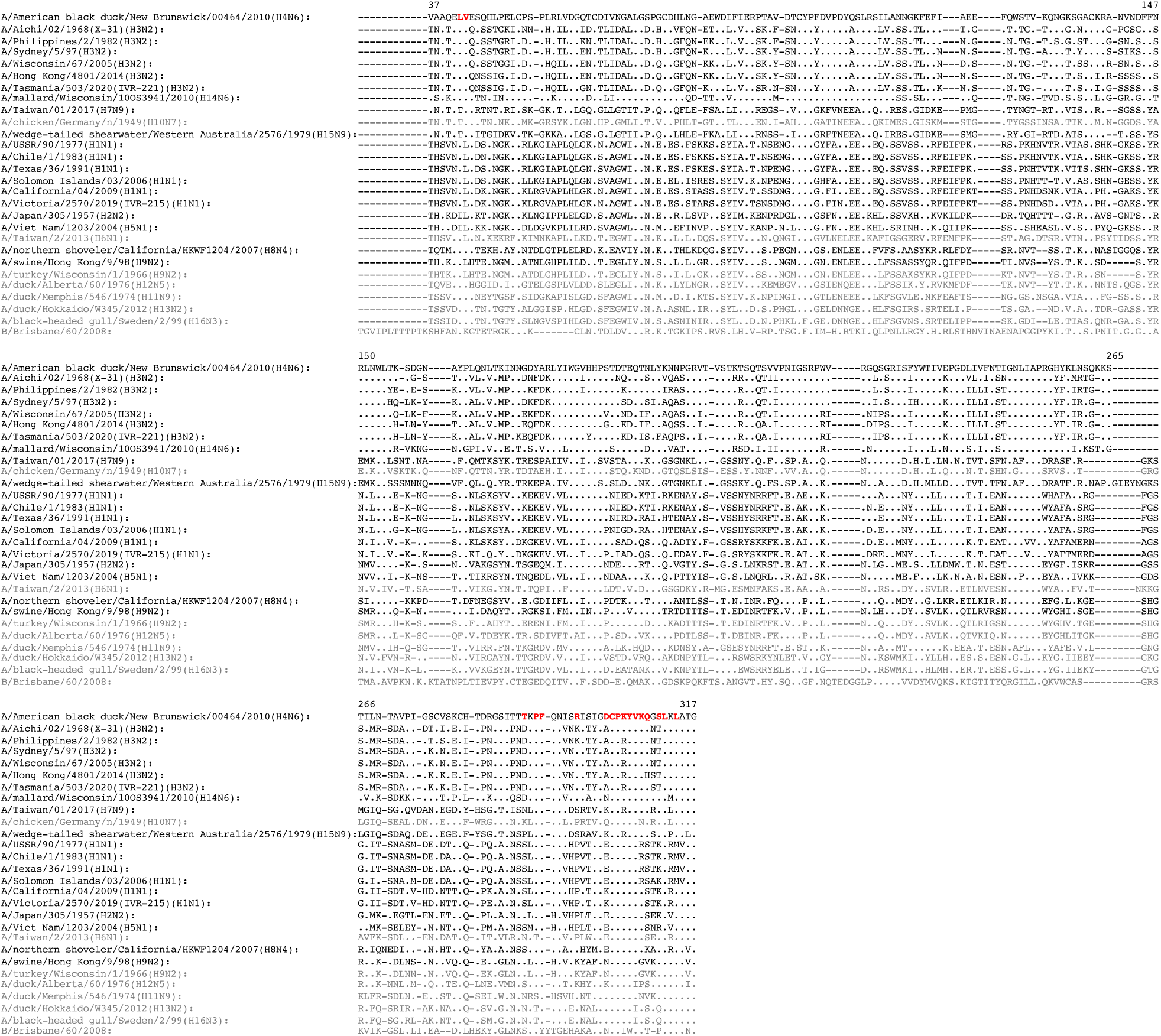
S8V1-157 Contacts conserved sites. An alignment of HA head regions for HAs used in this study. S8V1-157 contacting residues in A/American black duck/New Brunswick/00464/2010(H4N6) are colored red. HAs not bound by S8V1-157 are in denoted in gray. Sites identical to the A/American black duck/New Brunswick/00464/2010(H4N6) sequence are indicated by a “.”. Amino acid numbering corresponds to standard H3N2 numbering.

**Figure S2:**
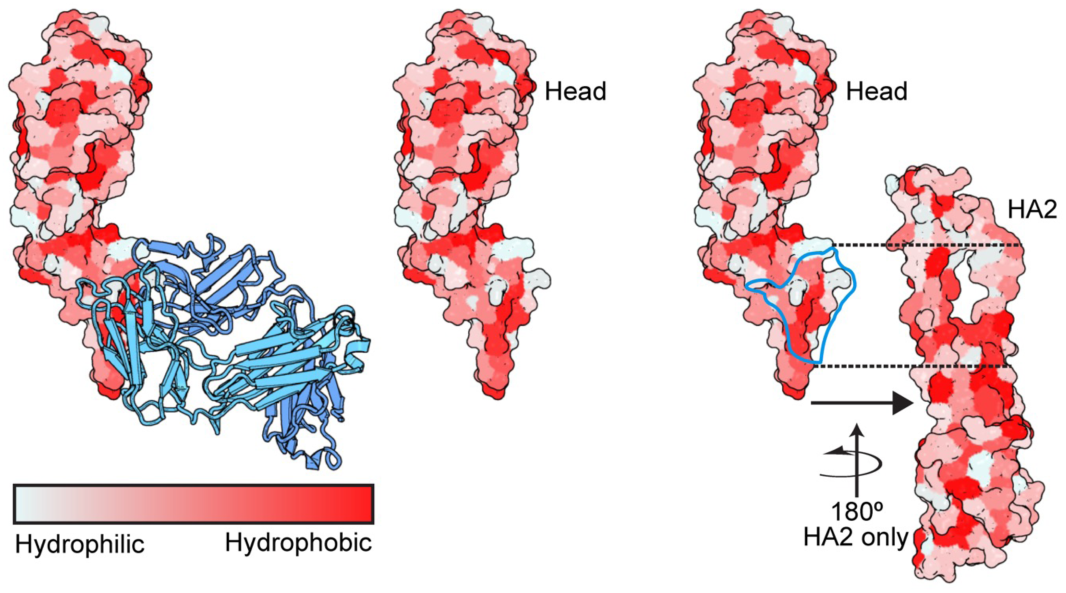
Surface hydrophobicity on the HA molecule. Surface hydrophobicity and the head-stem epitope are shown on the HA trimer of A/American black duck/New Brunswick/00464/2010(H4N6) (PDB: 5XL2) ^21^. Coloring is based on the PyMOL Color h script that utilizes a normalized consensus hydrophobicity scale^62^. Views and orientations match those in Figure 2. The head-stem epitope is circled in the far right panel.

**Figure S3:**
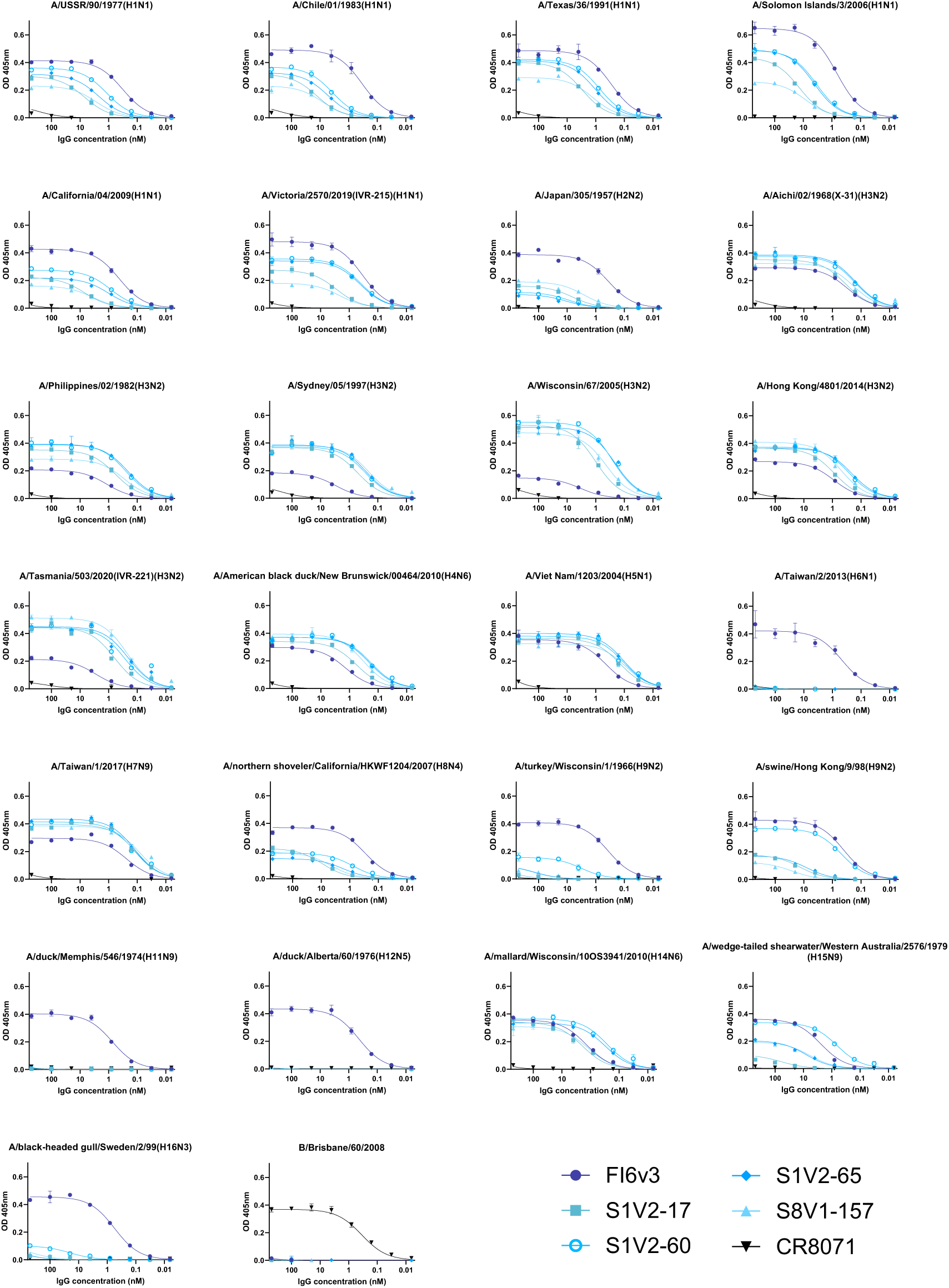
ELISA titrations of antibodies on HA coated plates. The broadly binding stem antibody FI6v3^47^ was used as a positive control and an influenza B specific head antibody, CR8071^58^, as a negative control for influenza A isolates. Data points represent the average of three technical replicates. The standard error of the mean is shown for each point. KDs were calculated from the curves fit to these data points.

**Figure S4:**
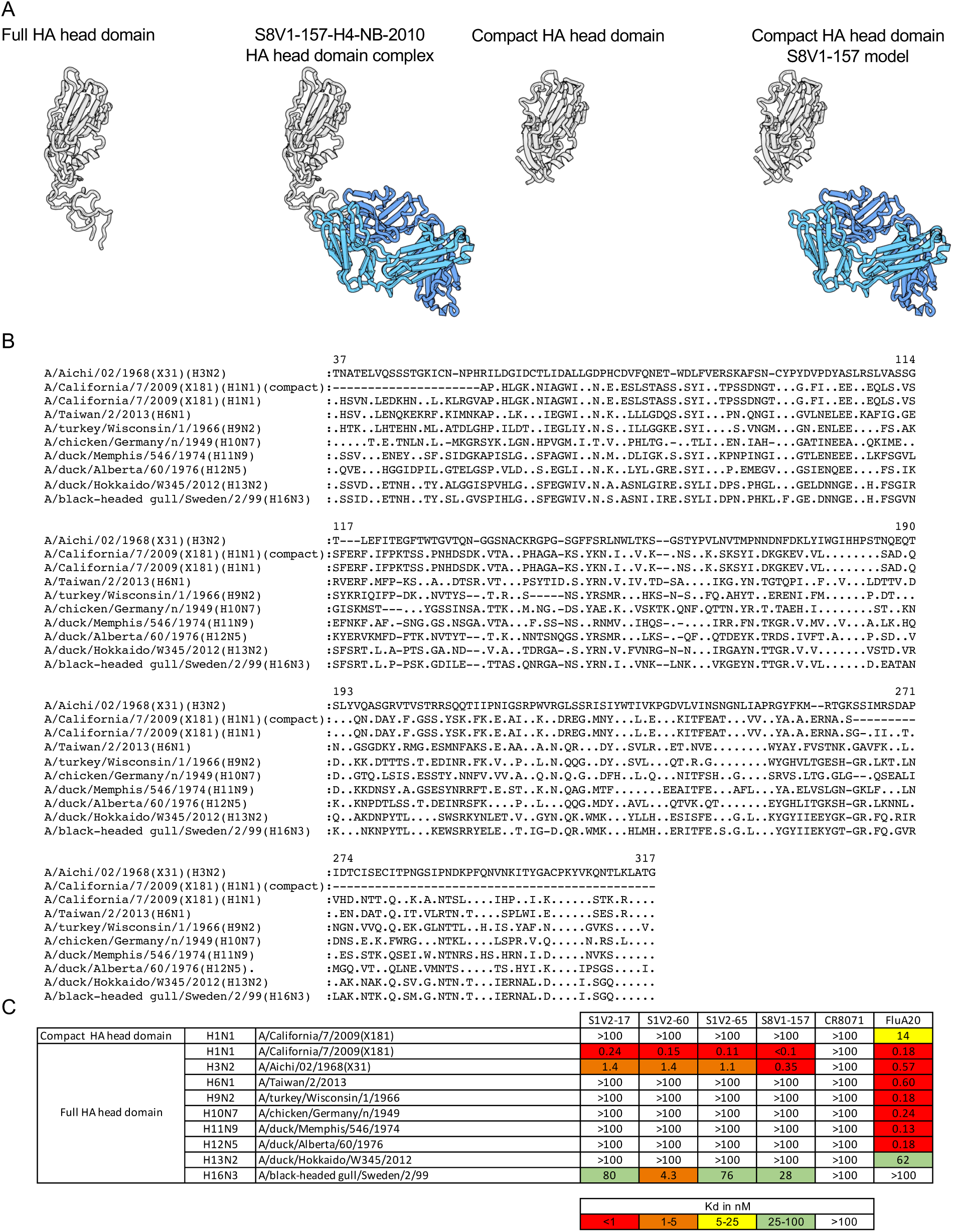
Head-stem epitope antibody binding to HA head-only constructs. A. Structures of HA head used in these experiments the S8V1-157-HA head complex is shown for reference. A compact head^49^, is shown for reference (PDB 7TRH). B. HA heads for HAs not bound or not expressed in Figure 3 were produced alongside a positive control A/California/07/2009(H1N1)(X-181) and a compact head version. Sequences of the HA head regions are aligned to the A/Aichi/02/1968(H3N2)(X-31) reference sequence and numbered by H3 convention. C. Dissociation constants from ELISA measurements of head-stem epitope antibodies to HA heads. Head interface antibody FluA-20^40^ was used as a positive control and influenza B head antibody CR8071^58^ as a negative control.

**Figure S5:**
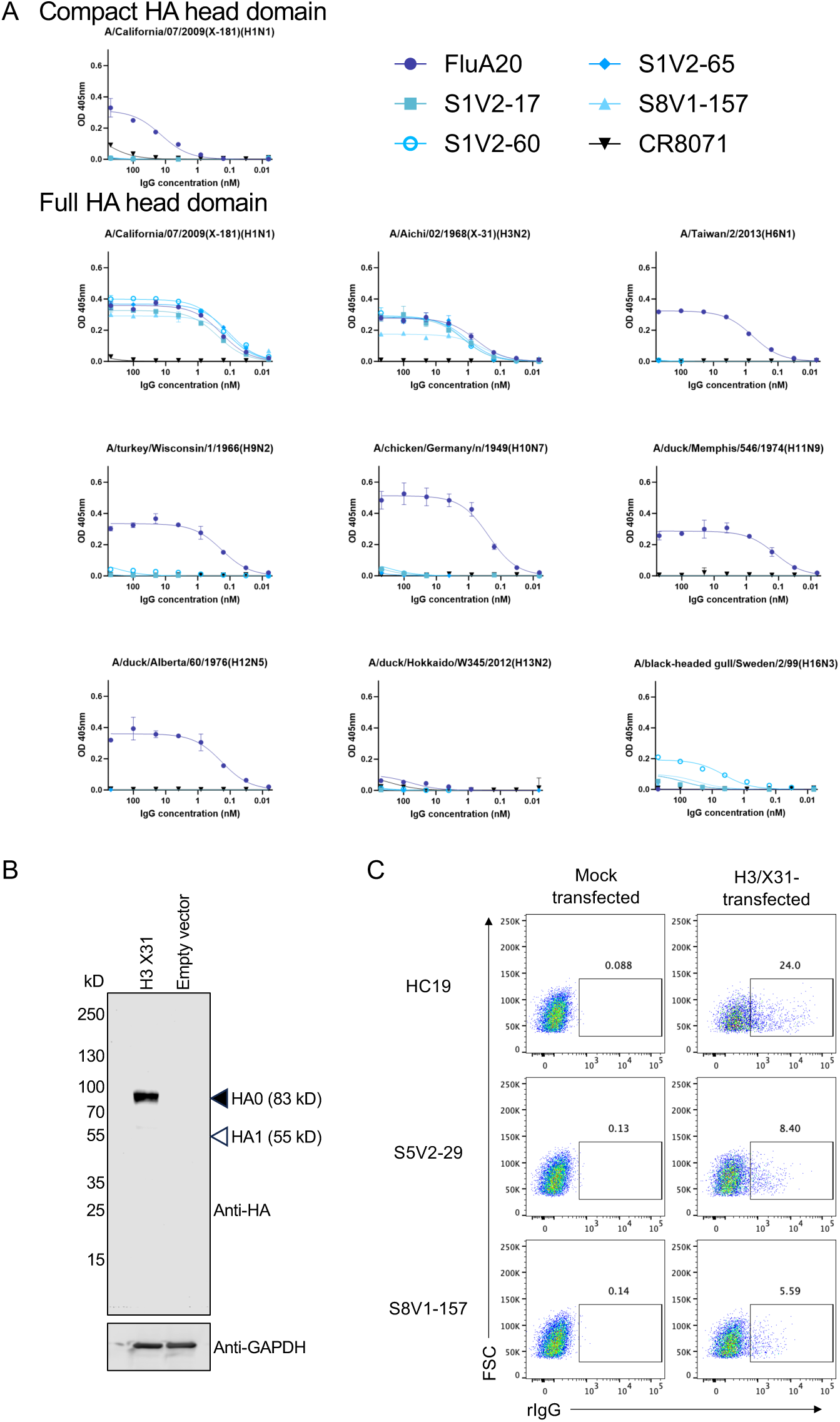
ELISA titrations of antibodies on HA head coated plates and characterization of cell expressed HA. A. Head interface antibody FluA-20^40^ was used as a positive control and influenza B head antibody CR8071^58^ as a negative control. Data points represent the average of three technical replicates. The standard error of the mean is shown for each point. KDs were calculated from the curves fit to these data points. B. Lysates from 293F transfected with either HA from A/Aichi/02/1968(H3N2)(X31) or empty vector were subjected to western blotting under reducing and denaturing conditions with either anti-HA tag antibody, which recognizes an endogenous sequence in the H3 HA head (HA1), or anti-GAPDH antibody. Molecular weights of unprocessed HA0 and processed HA1 are indicated with open or closed arrowheads, respectively. C. 293F transfected with either HA from A/Aichi/02/1968(H3N2)(X31) or empty vector were stained with the indicated antibodies and analyzed by flow cytometry. Gates denoting cells bound by antibody are shown, with the percent of the total population indicated above.

**Figure S6:**
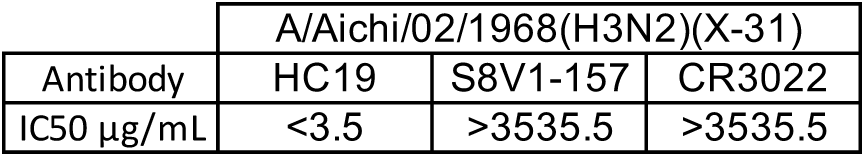
S8V1-157 is a non-neutralizing antibody. Neutralization IC50 values for S8V1-157, neutralizing antibody HC19^32^ (positive control) and SARS-CoV antibody CR3022^33^ (negative control).

**Figure S7:**
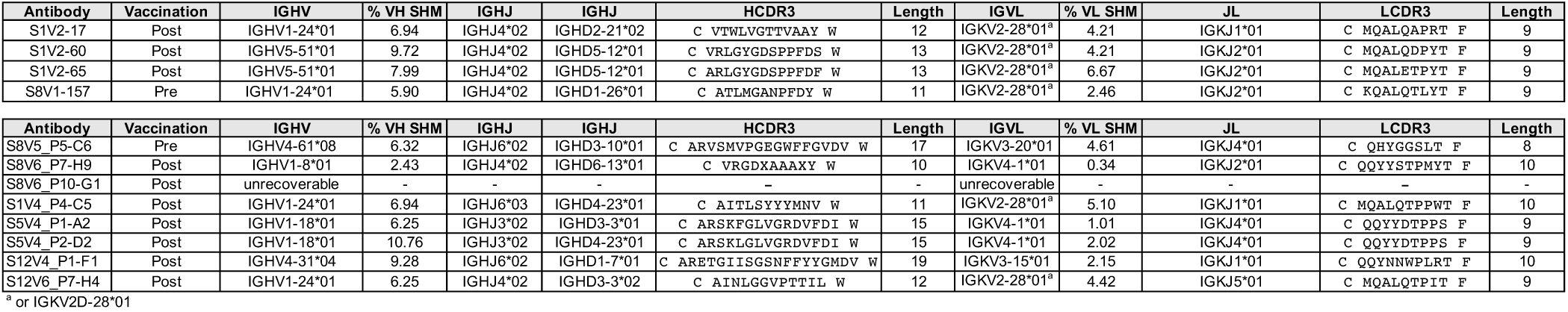
Antibody genetics of S8V1-157 competing antibodies. Antibody names, gene usage and HCDR3 sequences for S8V1-157 competing antibodies identified in Figure 6.

